# Differences in splicing defects between the grey and white matter in myotonic dystrophy type 1

**DOI:** 10.1101/819433

**Authors:** Masamitsu Nishi, Takashi Kimura, Mitsuru Furuta, Koichi Suenaga, Tsuyoshi Matsumura, Harutoshi Fujimura, Kenji Jinnai, Hiroo Yoshikawa

## Abstract

Myotonic dystrophy type 1 (DM1) is a multi-system disorder caused by CTG repeats in the *myotonic dystrophy protein kinase* (*DMPK*) gene. This leads to sequestration of the splicing factor, muscleblind-like 2 (MBNL2), and aberrant splicing, mainly in the central nervous system. We investigated the splicing patterns of *MBNL1/2* and genes controlled by MBNL2 in several regions of the brain and between the grey matter (GM) and white matter (WM) in DM1 patients using RT-PCR. Compared with the control, the percentage of spliced-in parameter (PSI) for most of the examined exons were significantly altered in most of the brain regions of DM1 patients, except for the cerebellum. The splicing of many genes was differently regulated between the GM and WM in both DM1 and control. The level of change in PSI between DM1 and control was higher in the GM than in the WM. The differences in alternative splicing between the GM and WM may be related to the effect of DM1 on the WM of the brain. We hypothesize that in DM1, aberrantly spliced isoforms in the neuronal cell body of the GM may not be transported to the axon. This might affect the WM as a consequence of Wallerian degeneration secondary to cell body damage. Our findings may have implications for analysis of the pathological mechanisms and exploring potential therapeutic targets.

## Introduction

Myotonic dystrophy type 1 (DM1) is the most common form of muscular dystrophy in adults, affecting the skeletal muscle, heart, ocular lens, testis, and central nervous system (CNS). The CNS symptoms of DM1 can have a negative impact on patient quality of life [1]. DM1 is caused by the unstable expansion of CTG trinucleotide repeats in the 3’ untranslated region (UTR) of the *myotonic dystrophy protein kinase* (*DMPK*) gene. These CTG repeats are transcribed to CUG repeats, leading to the formation of an RNA hairpin loop. These loops form foci in the nucleus and sequester splicing factors, such as the muscleblind-like (MBNL) protein. The MBNL protein controls splicing; in the nucleus, MBNL binds to the YGCY motif of pre-mRNAs and controls the splicing of alternative exons in either an exon inclusion or exon exclusion direction, depending on whether the motif is located in the downstream or upstream intron for each exon, respectively [2]. We have previously detected several splicing defects in the brains of both *MBNL1/2* knockout mice and DM1 patients, and revealed that MBNL2 acts as major splicing factor in the brain, while MBNL1 performs a similar function in the skeletal muscle [3, 4]. In addition, conditional double-knockout *Mbnl1* and *Mbnl2* mice showed greater splicing defects than either *Mbnl1* or *Mbnl2* single knockout mice, indicating that the combined loss of MBNL1 and MBNL2 is necessary to recapitulate mis-splicing, since MBNL1 and MBNL2 compensate for each other [5, 6].

We previously examined the mis-splicing in DM1 patients for each area of the brain and found that mis-splicing in the cerebellum is less apparent than in any other area [3]. We also demonstrated the different degrees of mis-splicing that occurs among cell layers of the cerebellum [7]. These results clearly showed the heterogenicity of the brain and show that detailed splicing analysis for each cell or region is essential to clarify the pathomechanisms of this disorder.

It has been reported that fetal spliced isoforms increase in adult DM1 tissues as a result of MBNL1/2 deletion, since MBNL1/2 regulates the splicing switch from fetal to adult [4, 8-11]. *MBNL2* splicing is developmentally regulated; exon 5 (54nt.) and exon 8 (95nt.) are included in the fetal brain, while both exons are excluded in the adult brain [12].

Using RNA-sequencing, Mills et al. showed that the grey matter (GM) and white matter (WM) have distinct transcriptome profiles, including for alternative splicing (AS) [13]. Neuroimaging analysis in patients with DM1 revealed various changes with conventional MRI [14, 15], voxel-based morphometry (VBM) [16, 17], and diffusion-weighted imaging (DWI) tensor analysis [18-21]. Overall, the degree of change is greater in the WM than in the GM [17, 22].

Aberrant splicing of *MBNL1* in the DM1 brain has been previously reported [23], but that of *MBNL2* is still unknown. Considering distinct regions affected, as shown by neuroimaging, and the differences in splicing regulation between the GM and WM, it is very important determine how AS is controlled in GM and WM of a DM1 brain. However, little is known about this issue. Therefore, in this study, we examined the splicing patterns of *MBNL1/2*, and the other genes that controlled by MBNL2, among several brain regions (frontal and temporal lobes, hippocampus, the cerebellum), and between the GM and WM in DM1 patients.

## Materials and methods

### Ethics and written informed consent

This research was approved by the Ethics Committee of Hyogo College of Medicine (approval number: 93), and written informed consent for specimen research was obtained from patients themselves or their family members during the autopsy.

### MBNL1 and 2 DNA sequence and primer design

We searched the DNA sequences for *MBNL1* and *MBNL2* using the GENETYX^®^ NCBI database. *MBNL1* has over 50 variants according to the presence or absence of exon 5, 6, 7, 8, and 9, or changes in the non-coding region. *MBNL2* has 10 splicing variants according to the presence or absence of exons 5, 7, and 8. Using NCBI Primer-BLAST, we designed two sets of primers for each spliced gene; exon 4 forward and exon 6 reverse primers for exon 5 splicing, and exon 6 forward and exon 9 reverse primers for exons 7 and 8 splicing. In addition, we examined splicing patterns of other genes controlled by MBNL2: *ADD1* exon 15, *CACNA1D* exon 11, *CLASP2* exon 23a and 23b, *CSCNK1D* exon 9, *KCNMA1* exon 27a, *TANC2* exon 22a, *GRIN1* exon 4, *MAPT* exon 2 and exon 12, using the primers used in our previous report [3, 7] (Tables 1 and 2).

**Table 1.**
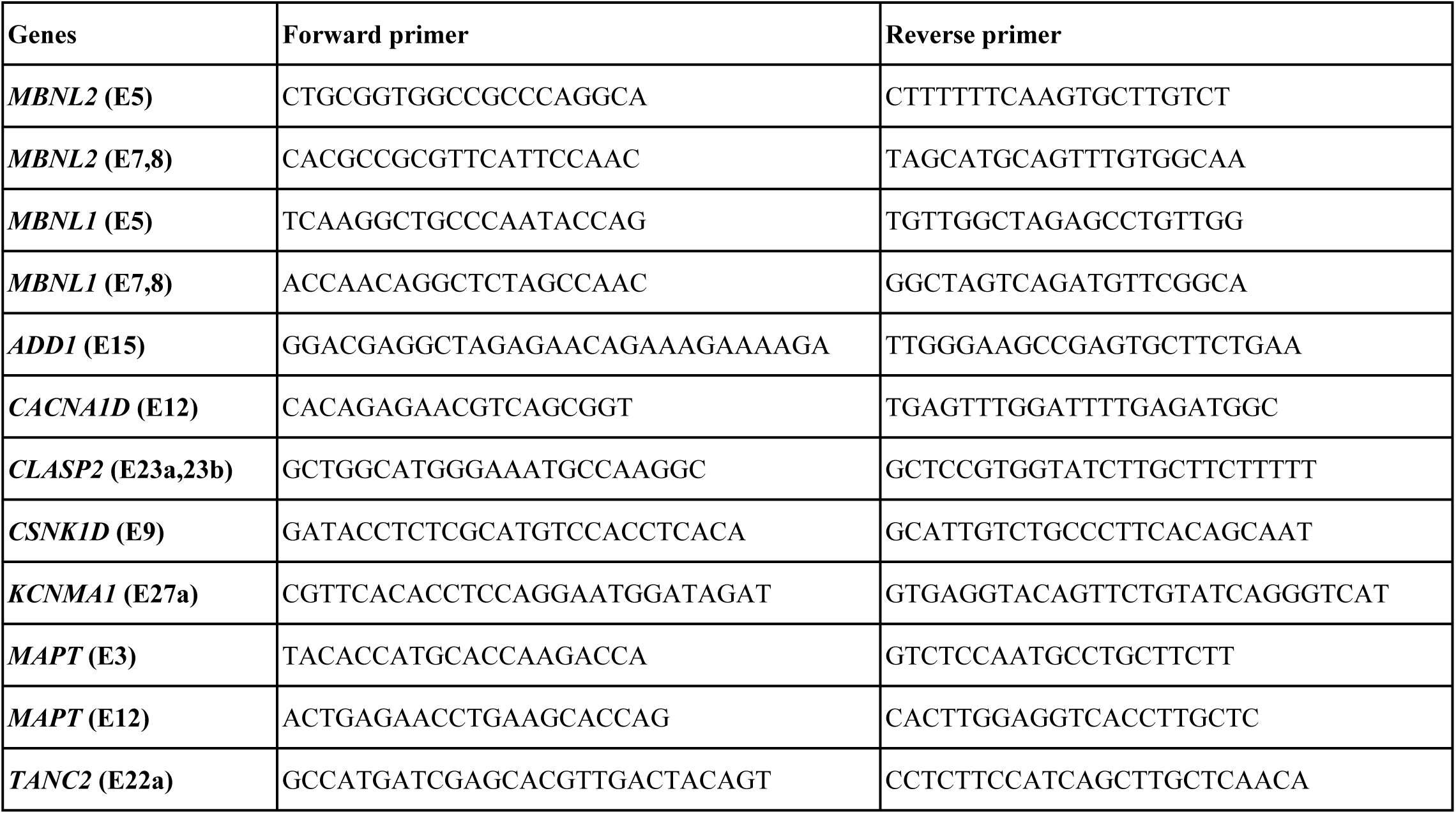
RT-PCR primers for alternative splicing.

**Table 2.**
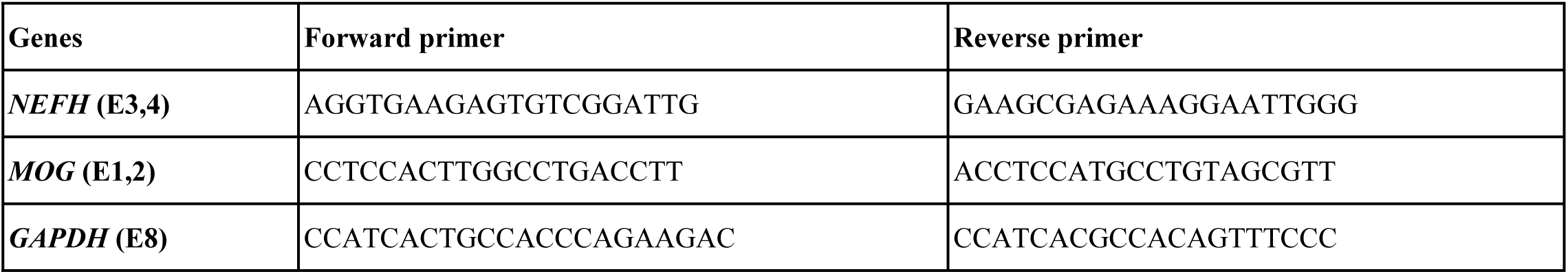
Quantitative RT-PCR primers for confirming the separation between the GM and WM

### Human RNA and splicing analysis

We investigated brains of patients during autopsy, with RNA extracted from 6 brains of patients with DM1 and 6 brains of patients with amyotrophic lateral sclerosis (as disease controls). Two RNA samples of fetal brains (as fetal controls) were obtained commercially (Cat. No. 540157, Agilent Technologies, US; Cat. No. 636526, Takara Bio Inc. Japan). We examined splicing patterns using comparison among several brain regions and between the GM and WM.

For comparison among several brain regions, RNA was extracted from the frontal and temporal lobes, the hippocampus, and the cerebellum using the ISOGEN^®^ reagent (Nippon Gene, Japan).

For comparison between the GM and WM, the frontal lobe tissue was sliced into several 20-μm thick sheets using a Cryostat (Leica^®^ CM1520, Germany). Before separation, one of the sheets was stained using the Luxol Fast Blue Stain Kit (ScyTek Laboratories Inc., US), to identify the boundary between the GM and WM. Only unstained sheets were used for RNA extraction. We separated each sheet into the GM and WM manually and extracted RNA using RNeasy^®^ Plus Mini (QIAGEN^®^, Germany) for comparison between the GM and WM.

To ensure separation of the GM and WM, we used the primers for *NEFH* (higher expression in the GM than the WM), *MOG* (higher expression in the WM than the GM), and *GAPDH* (control), and examined this by quantitative real-time PCR (qPCR) using the PowerUp™ SYBR™ with the 7500 Real-Time PCR system (Applied Biosystems, US). qPCR was performed in triplicates [13]. The levels of *NEFH* and *MOG* mRNA expression were calculated by the 2^-ΔΔCt^ method, using *GAPDH* as an endogenous control, and the data were presented as the mean of triplicates.

All cDNA was synthesized using the extracted RNA and purchased RNA (1 μg of RNA was used for comparison among several brain regions; 10-400 ng of RNA was used for comparison between the GM and WM) and random hexamers with the SuperScript^®^ III First-Strand Synthesis System (Invitrogen™, US) for reverse transcription PCR (RT-PCR) according to manufacturer’s instructions (Applied Biosystems). cDNA was amplified using AmpliTaq Gold^®^ 360 Master Mix (Applied Biosystems), with the initial denaturation set at 94°C for 10 min, followed by 36 (for comparison among several brain regions) or 42 (for comparison between the GM and WM) amplification cycles at 94°C for 30 s, 61°C for 30 s, and 72°C for 60 s. PCR products were analyzed with an Agilent 2100 Bioanalyzer (Agilent Technologies, US). We used percent spliced-in (PSI) values, indicating inclusion ratio of an alternative exon [3].

### Western Blotting

Similar to the comparison between the GM and WM, autopsied frontal lobe tissue sheets were separated into the GM and WM and homogenized using the RIPA Lysis Buffer System (ChemCruz, US). Equal amounts of protein (10 µg) were separated by SDS–PAGE and transferred onto Immune-Blot polyvinylidene difluoride membranes (Bio-Rad Laboratories, US). Blots were blocked with 5% (w/v) non-fat milk for 1 h and then incubated overnight at 4°C against the MBNL2 antibody (1:200, mouse monoclonal antibody raised against recombinant MBNL2 of human origin, sc-136167; Santa Cruz Biotechnology) or β-actin (1:1000, mouse polyclonal antibody against synthetic peptide corresponding to Human beta Actin aa 1-100 conjugated to keyhole limpet hemocyanin, ab8226; Abcam, UK). After repeated washings, the membranes were incubated at 4°C overnight with a peroxidase-conjugated anti-mouse immunoglobulin G (1:10,000, goat polyclonal antibody, 62-6520; Invitrogen). The membranes were then washed, developed using a chemiluminescence kit (GE Healthcare Life Sciences, US), and imaged using the ImageQuant LAS4000 system (GE Healthcare Life Sciences).

### Statistical analysis

Statistical analysis was performed using Welch’s t-test (two-tailed distribution). No adjustment for overall alpha error level in consideration of multiple comparison was performed to retain maximum power in this study, as suggested by Saville [24]. A p value less than 0.05 was considered statistically significant.

## Results

### Comparison among several brain regions

Compared with the control, PSI for *MBNL1* exon 5 were higher in all examined brain areas of DM1. The difference between the PSI for DM1 and the control for *MBNL1* exon 8 was not statistically significant (Fig 1, S1 Fig). The result of *MBNL1* exon 5 splicing was similar to a previous report [23], and that of exon 8 was not reported. Compared with the control, PSI for *MBNL2* exon 5 and exon 8 were significantly higher in most brain areas of DM1, except for the cerebellum. Notably, for the frontal region, temporal region, and hippocampus of DM1, PSI for *MBNL1* and *MBNL2* exons 5 and exon 8 were highly variable compared with the control.

**Figure 1.**
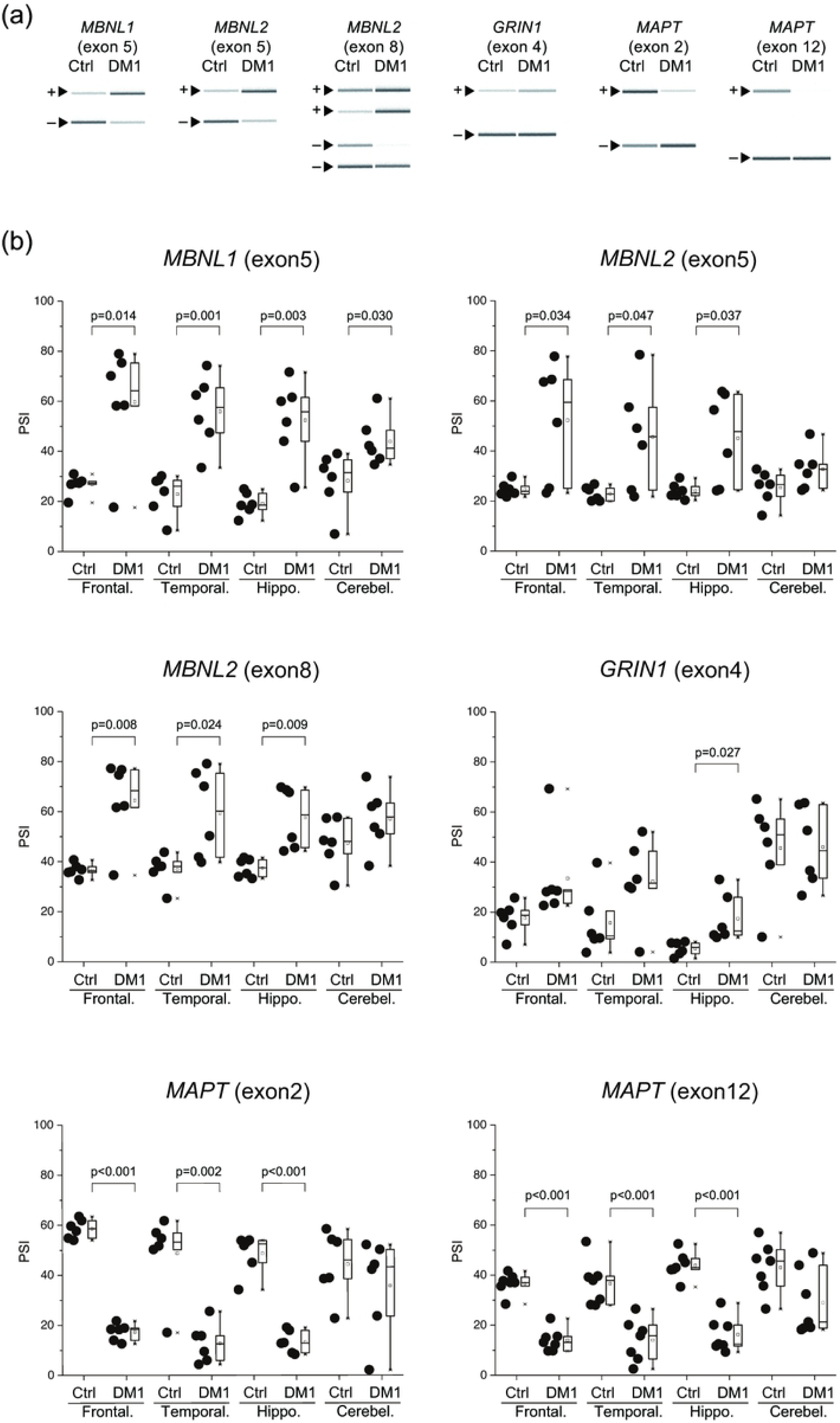
Aberrant splicing among several regions of the brain. (a) Representative RT-PCR products from the frontal lobe in the control and DM1. (b) Inclusion ratios of splicing changes in several brain regions. PSI values of all examined genes were compared by a pairwise Welch’s T-test. DM1, myotonic dystrophy type 1; Frontal., Frontal lobe; Temporal., Temporal lobe; Hippo., Hippocampus; Cerebel., Cerebellum.

Compared with the control, PSI for *MAPT* exon 2 and exon 12 were significantly lower in most brain areas of DM1, except for the cerebellum. PSI for *GRIN1* exon 4 was significantly higher in the DM1 hippocampus than in the control. Other areas had no significant difference. PSI was very low in the fetal brain tissue.

### Comparison between the GM and WM

Q-PCR analysis showed that the expression levels of *NEFH* was higher in the GM, while that of *MOG* was higher in the WM (Fig 2), confirming that the GM and WM were correctly separated.

**Figure 2.**
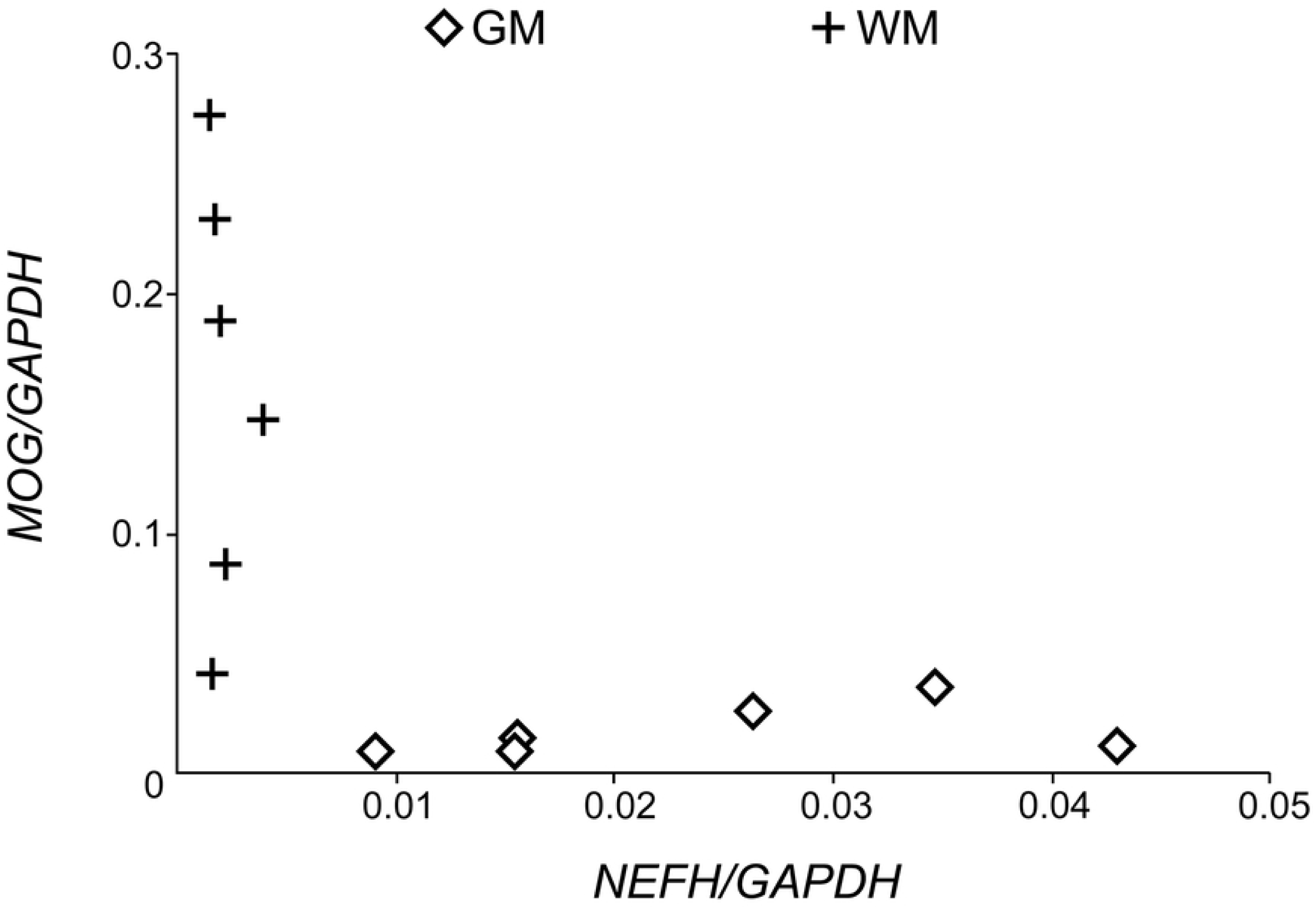
Expression level of *NEFH* and *MOG* in the GM and WM. Concentration of *NEFH* and *MOG* mRNA calculated by the 2^-ΔΔCt^ method, which was divided by that of *GAPDH*. Data are presented as the mean of triplicates. GM, grey matter; WM, white matter.

In the fetal brain, there were two patterns of AS: 1) exon including type (*ADD1, CLASP2, TANC2*), 2) exon excluding type (*MBNL1/2* exon 5 and exon 8, *CACNA1D, CSNK1D, KCNMA1, MAPT, GRIN1*) (Fig 3, S2 Fig).

**Figure 3.**
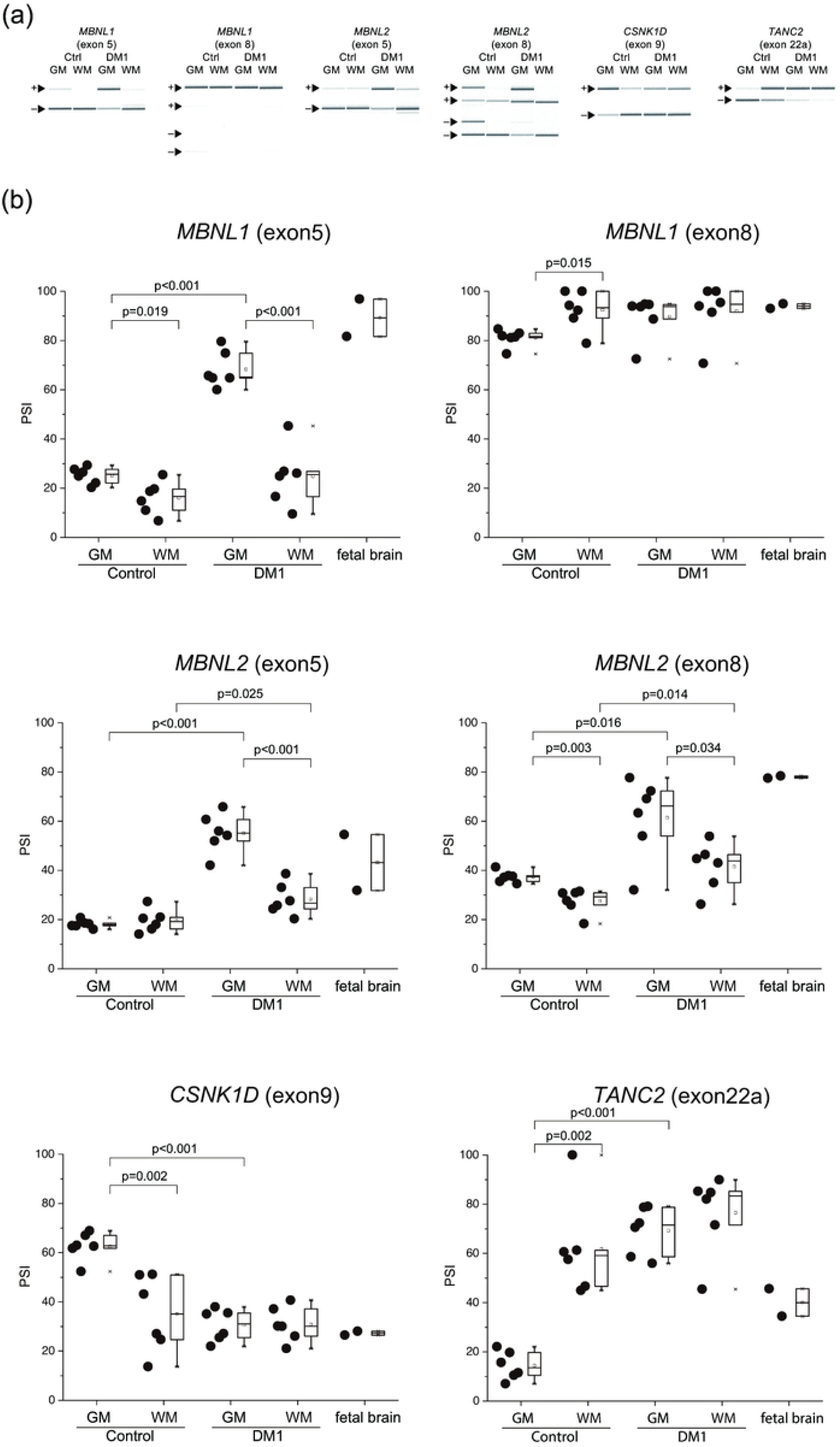
Aberrant splicing between the GM and WM. (a) Representative RT-PCR products from the frontal lobe GM and WM in control and DM1. (b) Inclusion ratios of splicing changes in the GM and WM. PSI values of all examined genes were compared by pairwise Welch’s T-test. GM, grey matter; WM, white matter.

In the control, PSI was significantly different between the GM and WM in 12 out of 13 AS events. Three types of AS were observed: 1) no change between the GM and WM (*GRIN1*), 2) increasing fetal splicing isoforms in the WM *(MBNL1* exon 8, *ADD1, CACNA1D, CSNK1D, TANC2, KCNMA1, MAPT* exon 2), 3) decreasing fetal splicing isoforms in the WM (*MBNL1* exon 5, *MBNL2* exon 5 and exon 8, *CLASP2, MAPT2* exon 12).

We calculated the level of PSI change (ΔPSI), which was obtained by subtracting the average of PSI between DM1 and the control. In most genes, ΔPSI of the GM was higher than ΔPSI of the WM, suggesting that more splicing misregulation occurs in the GM. For example, average PSI of *MBNL2* for exon 8 were 61.38%, 41.53%, 37.33%, and 27.50% in the GM and WM of DM1, and the GM and WM of control, respectively. ΔPSI of the GM was 24.05%, while ΔPSI of the WM was 14.03%.

PSI of *GRIN1* exon 4 showed no statistically significant difference between DM1 and the control, or between the GM and WM.

### Western Blotting

Western blotting analysis showed no significant differences in expression levels of MBNL2 protein between the GM and WM (Fig 4).

**Figure 4.**
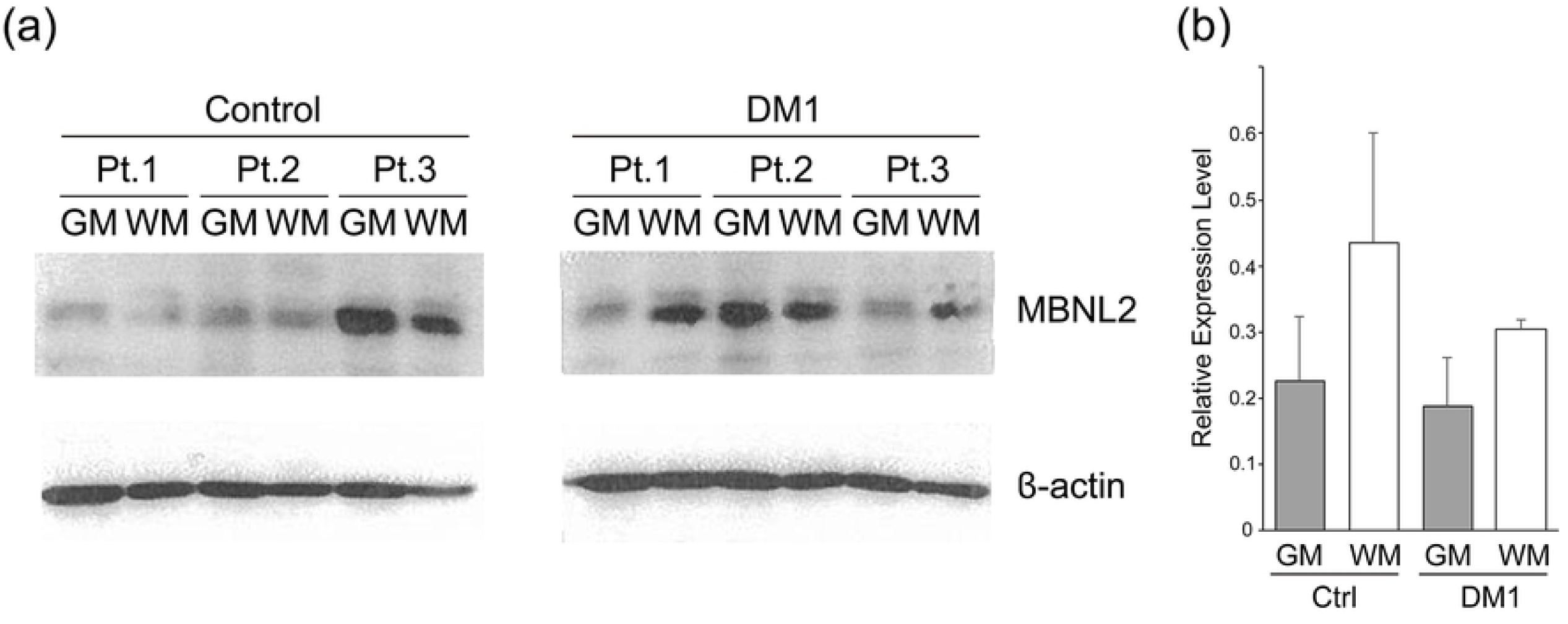
Expression of levels of MBNL2 and β-actin proteins. (a) Western blot of lysates from the GM and WM of DM1 and the control brain. (b) The relative expression levels of MBNL2. The levels of MBNL2 were divided by those of β-actin, and the ratios are shown. There was no statistically significance (n = 3) by Welch’s t-test. Error bar indicates differences in the mean ± SEM. GM, grey matter; WM, white matter.

## Discussion

Comparison among several brain regions revealed that PSI of almost of all examined gene exons were higher in most of the brain regions of DM1 patients than in those of the controls, except for the cerebellum. Comparison between the GM and WM revealed that splicing of many genes was differently regulated between the GM and WM. The extent of splicing between DM1 and the control was higher in the GM than the WM.

According to our previous research, there were fewer alternative splicing defects in the DM1 cerebellum than in other brain regions [3]. We showed that the inclusion of *MBNL1* exon 5 and *MBNL2* exon 5 and exon 8 was higher in most brain areas, except the cerebellum for *MBNL2* exon 5 and exon 8. *MBNL1* exon 5 inclusion in the DM1 brain had previously been reported [23]; however, we are the first to demonstrate *MBNL2* exon 5 and exon 8 inclusion in the DM1 brain. Comparison of *MBNL1/2* among several brain regions was similar to those of previous reports [3], in that aberrant splicing is observed in most brain areas except for the cerebellum.

*MAPT* [25] and *GRIN1* [10] splicing defects were found in DM1 patient brains. The aberrant splicing of *MAPT* exon 2 and exon 3 [4], *MAPT* exon 10 and *GRIN1* [5] splicing was recapitulated by using knock out mice of *Mbnl1/2*, suggesting that these splicing events are controlled by the MBNL 1/2 protein. We analyzed both these genes in several areas of the brain and found that *MAPT* splicing was similar to that of other genes; there were fewer splicing changes in the cerebellum than in other brain areas in DM1. *GRIN1* was different in that: 1) In the hippocampus, PSI of DM1 was significantly higher than that of control. In the frontal and temporal lobes, PSI of DM1 tended to be higher than that of the control, but these differences did not reach statistical significance; 2) PSI in the fetal brain was lower than that in the adult control, in contrast to DM1. A significant difference was observed in the hippocampus between the control and DM1, but not in other areas. Since *GRIN1* had large variations in PSI values compared to the others, this difference may have caused difficulty in reaching significance.

Analysis between the GM and WM revealed that, compared to the control, splicing changes of DM1 in the GM exceeded those of WM, except for *GRIN1*. In the Comparison among several brain regions, we collected RNA from both the GM and WM, and did not evaluate the ratio of GM to WM. Therefore, we assume that the distribution of PSI observed in the comparison among several brain regions may have been caused by the variation in the ratio of GM/WM in the collected samples.

Western blot analysis demonstrated only one band for MBNL2, and there was no significant difference between DM1 and control or between the GM and WM. Including exon 5 of *MBNL2*, the molecular mass of MBNL2 protein increases by about 2 kDa. Including exon 8 of *MBNL2*, the molecular mass does not change significantly. This is because the exon 8 inclusive isoform 1 protein contains 367 amino acids, while exon 8 exclusive, but otherwise identical, isoform 4 contains 373 amino acids. Therefore, we assumed that there may be some MBNL2 bands for several spliced isoforms, with or without exon 5 and exon 8. Antibodies recognizing the peptides corresponding to exon 5 and exon 8 would be useful to examine the expression of each isoform.

This study revealed that the degree of mis-splicing in DM1 differs between the GM and WM. The GM is composed of the neuronal cell bodies, protoplasmic astrocytes, and microglial cells, while the WM comprises axons, oligodendrocytes, fibrous astrocytes, microglial cells, and ependymal cells (near the brain ventricle) [26]. In this study, as mRNA was extracted from the WM just under the GM, ependymal cells were not included in the samples. There are a few possibilities that could explain why these differences in splicing between the GM and WM occurred. One possibility is that these differences may be affected by the distribution of each splicing isoform between the axon and neuronal cell body. The WM is less transcriptionally active than the GM, and mRNA in the WM may represent mRNA flow from the GM [13], since the mRNAs are transcribed in the neuronal cell body, and part of them are transported to the axon [27]. *MBNL1/2* exon 5 includes nuclear localization signals, and it is reported that the isoforms with this exon tend to be exclusively expressed in the nucleus [12]. Inclusion of exon 8 reduces the hairpin loop mobility of MBNL2 [12]. Since both exons tend to be present in DM1 patients, fewer free MBNL proteins in the cytoplasm should be present. In order to target to specific compartments, MBNL binds to the distal 3’UTR protein binding sites and may intercede isoform specific mRNA localization [28]. Hence, it is possible that presence or absence of each alternative exon may influence its binding ability to the 3’UTR of *MBNL* and the transport of its mRNA. Consequently, aberrant splicing of *MBNL* mRNA and resulting intracellular localization shift of MBNL in DM1 patients might affect axonal transport of each spliced mRNA isoform. Another possibility is that splicing differences between the GM and WM may be influenced by splicing patterns of other cells such as oligodendrocytes, fibrous astrocytes, and microglial cells of the WM. These distinct splicing changes may be due to differences in the number of CTG repeats in the cells between the GM and WM. Jinnai et al. examined the somatic instability of the CTG expansion in various regions of the CNS and found that the expansion in the GM was longer than that in the WM, although this difference did not reach statistical significance [29]. Possibly, this relatively short expansion in the WM might explain the modest level of aberrant splicing. In order to examine this possibility, it is necessary to isolate each cell using cell culture or laser microdissection.

VBM and DWI tensor studies have shown that the effect of DM1 on the WM is more prominent than on the GM and that WM T2 hyperintensity lesions were found in the frontal, temporal (especially at the anterior temporal poles), and parietal lobes [21, 22]. However, this study showed more splicing defects in the GM than in the WM. It is unclear how these differences in the degree of splicing abnormalities relate to the predominant WM change. It remains debated whether the change in the WM is a result of Wallerian degeneration or a primary process of DM1 [17, 19]. As mentioned above, we suggested two possibilities explaining the differences in splicing defects between the GM and WM. Assuming the possibility that aberrant/fetal splicing mRNA isoforms are difficult to transport to the axon, aberrantly spliced isoforms would increase in the neuronal cell body, which could cause axonal injury as a consequence of Wallerian degeneration. The possibility that fewer splicing defects occurred in the myelin sheath of oligodendrocytes compared to the neuronal cell body does not explain the predominantly affected WM as shown by neuroimaging. Taken together, we hypothesize that WM lesions were caused by Wallerian degeneration due to neuronal cell body damage. Although there are limited findings on how each splicing isoform protein is distributed, this is the first study showing the differences in splicing regulation and the degree of aberrant splicing in DM1 between the GM and WM.

## Conclusion

In this study, we showed that splicing changes are relatively modest in the WM compared to the GM of the DM1 brain, which indicated the possibility that aberrant/fetal splicing isoforms may not be transported to the axon. Our result suggests that the predominant effects of DM1 on the WM might represent axonal injury by Wallerian degeneration due to GM dominant aberrant splicing. We believe that exploring the distribution of each spliced protein isoform by immunostaining can help in our understanding of the pathological mechanisms of DM1. Future studies should also investigate mRNA and protein transport and local translation in the axons of neuronal cells.

## Acknowledgements

The authors thank the Research Resource network Japan for the human brain samples.

## Conflicts of interest

There are no conflicts of interest.

## Supporting information

**S1 Fig. MBNL1 exon 8 aberrant splicing among several regions of the brain.**

(a) Representative RT-PCR products from the frontal lobe in the control and DM1. (b) Inclusion ratios of splicing changes in several brain regions. PSI values of MBNL1 exon 8 was compared by a pairwise Welch’s T-test. DM1, myotonic dystrophy type 1; Frontal., Frontal lobe; Temporal., Temporal lobe; Hippo., Hippocampus; Cerebel., Cerebellum.

**S2 Fig. Aberrant splicing between the GM and WM.**

(a) Representative RT-PCR products from the frontal lobe GM and WM in control and DM1. (b) Inclusion ratios of splicing changes in the GM and WM. PSI values of all examined genes were compared by pairwise Welch’s T-test. GM, grey matter; WM, white matter.

## References

1. Bugiardini E, Meola G, Group D-C. Consensus on cerebral involvement in myotonic dystrophy: workshop report: May 24-27, 2013, Ferrere (AT), Italy. Neuromuscul Disord. 2014;24(5):445–52.

2. Zhang C, Lee KY, Swanson MS, Darnell RB. Prediction of clustered RNA-binding protein motif sites in the mammalian genome. Nucleic Acids Res. 2013;41(14):6793–807.

3. Suenaga K, Lee KY, Nakamori M, Tatsumi Y, Takahashi MP, Fujimura H, et al. Muscleblind-like 1 knockout mice reveal novel splicing defects in the myotonic dystrophy brain. PLoS One. 2012;7(3):e33218.

4. Charizanis K, Lee KY, Batra R, Goodwin M, Zhang C, Yuan Y, et al. Muscleblind-like 2-mediated alternative splicing in the developing brain and dysregulation in myotonic dystrophy. Neuron. 2012;75(3):437–50.

5. Goodwin M, Mohan A, Batra R, Lee KY, Charizanis K, Fernández Gómez FJ, et al. MBNL Sequestration by Toxic RNAs and RNA Mispro-cessing in the Myotonic Dystrophy Brain. Cell Rep. 2015;12(7):1159–68.

6. Lee KY, Li M, Manchanda M, Batra R, Charizanis K, Mohan A, et al. Compound loss of muscleblind-like function in myotonic dystrophy. EMBO Mol Med. 2013;5(12):1887–900.

7. Furuta M, Kimura T, Nakamori M, Matsumura T, Fujimura H, Jinnai K, et al. Macroscopic and microscopic diversity of missplicing in the central nervous system of patients with myotonic dystrophy type 1. Neuroreport. 2018;29(3):235–40.

8. Braz SO, Acquaire J, Gourdon G, Gomes-Pereira M. Advances in the Understanding of Neuromuscular Aspects of Myotonic Dystrophy. Front Neurol. 2018;9:519.

9. Lin X, Miller JW, Mankodi A, Kanadia RN, Yuan Y, Moxley RT, et al. Failure of MBNL1-dependent post-natal splicing transitions in myo-tonic dystrophy. Hum Mol Genet. 2006;15(13):2087–97.

10. Jiang H, Mankodi A, Swanson MS, Moxley RT, Thornton CA. Myotonic dystrophy type 1 is associated with nuclear foci of mutant RNA, sequestration of muscleblind proteins and deregulated alternative splicing in neurons. Hum Mol Genet. 2004;13(24):3079–88.

11. Kanadia RN, Johnstone KA, Mankodi A, Lungu C, Thornton CA, Esson D, et al. A muscleblind knockout model for myotonic dystrophy. Science. 2003;302(5652):1978–80.

12. Sznajder Ł, Michalak M, Taylor K, Cywoniuk P, Kabza M, Wojtkowiak-Szlachcic A, et al. Mechanistic determinants of MBNL activi-ty. Nucleic Acids Res. 2016;44(21):10326–42.

13. Mills JD, Kavanagh T, Kim WS, Chen BJ, Kawahara Y, Halliday GM, et al. Unique transcriptome patterns of the white and grey matter cor-roborate structural and functional heterogeneity in the human frontal lobe. PLoS One. 2013;8(10):e78480.

14. Abe K, Fujimura H, Toyooka K, Yorifuji S, Nishikawa Y, Hazama T, et al. Involvement of the central nervous system in myotonic dystrophy. J Neurol Sci. 1994;127(2):179–85.

15. Ogata A, Terae S, Fujita M, Tashiro K. Anterior temporal white matter lesions in myotonic dystrophy with intellectual impairment: an MRI and neuropathological study. Neuroradiology. 1998;40(7):411–5.

16. Schneider-Gold C, Bellenberg B, Prehn C, Krogias C, Schneider R, Klein J, et al. Cortical and Subcortical Grey and White Matter Atrophy in Myotonic Dystrophies Type 1 and 2 Is Associated with Cognitive Im-pairment, Depression and Daytime Sleepiness. PLoS One. 2015;10(6):e0130352.

17. Minnerop M, Weber B, Schoene-Bake JC, Roeske S, Mirbach S, Anspach C, et al. The brain in myotonic dystrophy 1 and 2: evidence for a predominant white matter disease. Brain. 2011;134(Pt 12):3530–46.

18. Fukuda H, Horiguchi J, Ono C, Ohshita T, Takaba J, Ito K. Diffusion tensor imaging of cerebral white matter in patients with myotonic dystrophy. Acta Radiol. 2005;46(1):104–9.

19. Ota M, Sato N, Ohya Y, Aoki Y, Mizukami K, Mori T, et al. Relationship between diffusion tensor imaging and brain morphology in pa-tients with myotonic dystrophy. Neurosci Lett. 2006;407(3):234–9.

20. Wozniak JR, Mueller BA, Ward EE, Lim KO, Day JW. White matter abnormalities and neurocognitive correlates in children and adolescents with myotonic dystrophy type 1: a diffusion tensor imaging study. Neuromuscul Disord. 2011;21(2):89–96.

21. Okkersen K, Monckton DG, L. N, Tuladhar AM, Raaphorst J, van Engelen BGM. Brain imaging in myotonic dystrophy type 1: A systematic review. Neurology. 2017;89(9):960–9.

22. Minnerop M, Gliem C, Kornblum C. Current Progress in CNS Imaging of Myotonic Dystrophy. Front Neurol. 2018;9:646.

23. Dhaenens CM, Schraen-Maschke S, Tran H, Vingtdeux V, Ghanem D, Leroy O, et al. Overexpression of MBNL1 fetal isoforms and modified splicing of Tau in the DM1 brain: two individual consequences of CUG trinucleotide repeats. Exp Neurol. 2008;210(2):467–78.

24. Saville DJ. Multiple Comparison Procedures: The Practical Solution. The American Statistician. 1990;44(2):174–80.

25. Sergeant N, Sablonnière B, Schraen-Maschke S, Ghestem A, Maurage CA, Wattez A, et al. Dysregulation of human brain microtubule-associated tau mRNA maturation in myotonic dystrophy type 1. Hum Mol Genet. 2001;10(19):2143–55.

26. Mescher AL, Junqueira LCU. Junqueira’s basic histology: texts and atlas. Fifteenth edition. ed. New York: McGraw Hill Education; 2018. ix, 562 pages p.

27. Cioni JM, Koppers M, Holt CE. Molecular control of local translation in axon development and maintenance. Curr Opin Neurobiol. 2018;51:86–94.

28. Wang ET, Cody NA, Jog S, Biancolella M, Wang TT, Treacy DJ, et al. Transcriptome-wide regulation of pre-mRNA splicing and mRNA lo-calization by muscleblind proteins. Cell. 2012;150(4):710–24.

29. Jinnai K, Mitani M, Futamura N, Kawamoto K, Funakawa I, Itoh K. Somatic instability of CTG repeats in the cerebellum of myotonic dys-trophy type 1. Muscle Nerve. 2013;48(1):105–8.

